# A New CUT&RUN Low Volume-Urea (LoV-U) protocol uncovers Wnt/β-catenin tissue-specific genomic targets

**DOI:** 10.1101/2022.07.06.498999

**Authors:** Gianluca Zambanini, Anna Nordin, Mattias Jonasson, Pierfrancesco Pagella, Claudio Cantù

## Abstract

Upon WNT/β-catenin pathway activation, stabilized β-catenin travels to the nucleus where it associates with the TCF/LEF family of transcription factors, which constitutively bind to genomic Wnt Responsive Elements (WREs), to activate transcription of target genes. Discovering the binding profile of β-catenin is therefore required to unambiguously assign direct targets of WNT signaling. Cleavage Under Target and Release Using Nuclease (CUT&RUN) has recently emerged as a prime technique for mapping the binding profile of chromatin interacting proteins. In our attempts to profile different regulators of the WNT/β-catenin transcriptional complex, CUT&RUN performed reliably when targeting transcription factors such as TCF/LEF, but it failed to produce consistent binding patterns of the non-DNA-binding β-catenin. Here, we present a biochemical modification of the CUT&RUN protocol, which we refer to as LoV-U (Low Volume and Urea), that enables the generation of robust and reproducible β-catenin binding profiles. CUT&RUN-LoV-U uncovers direct WNT/β-catenin target genes in human cells, as well as in ex *vivo* cells isolated from developing mouse tissue. CUT&RUN-LoV-U can profile all classes of chromatin regulators tested and is well suited for simultaneous processing of several samples. We submit that the application of our protocol will allow the detection of the complex system of tissue-specific WNT/β-catenin target genes, together with other non-DNA-binding transcriptional regulators that act downstream of ontogenetically fundamental signaling cascades.

## Introduction

Gene regulation is achieved by a combinatorial system of DNA-binding transcription factors (TFs) which physically associate with specific DNA sequences within regulatory regions in the genome, and non-DNA-binding co-factors (Co-Fs) which are recruited by TFs or histone marks and serve as hubs to tether chromatin modifying complexes and RNA Polymerase II (Klemm et al., 2019). The molecular apparatus necessary for gene transcription is therefore constituted by a complex assembly of several proteins that recruit each other to the DNA, whilst not necessarily binding to it (Mitsis et al., 2020; Nikolov and Burley, 1997). Characterizing the genome-wide binding profile and positioning of all these classes of chromatin interactors is crucial to understand the complexities of gene regulation and its dynamics (Merika and Thanos, 2001).

This is emphasized, in the WNT/β-catenin signaling pathway, an evolutionarily conserved intracellular cascade where the extracellular signal carried by WNT ligands is transduced in the signal-receiving cell into a gene expression program (Rim et al., 2022). Here, β-catenin serves as the pivotal protein for signal transduction from the cytosol into the nucleus, where it coordinates a transcriptional response (Valenta et al., 2012). By its nature and structure, β-catenin mediates protein-protein interactions, but it is not capable of directly binding to DNA (Huber et al., 1997). The locus-specific interaction of β-catenin is conferred by its physical association with the TCF/LEF family of TFs, which bind a consensus sequence found within WNT Responsive Elements (WRE) (Mosimann et al., 2009). Subsequently, β-catenin recruits a series of Co-Fs, such as BCL9/BCL9L, PYGO1/2 (Kramps et al., 2002), and CBP/p300 (Takemaru and Moon, 2000), to engage the basic machinery of transcription and activate target genes (Söderholm and Cantù, 2020). As a consequence, chromatin-associated β-catenin becomes embedded within a large transcriptional complex, sometimes referred to as the WNT enhanceosome (van Tienen et al., 2017). The detection of the chromatin binding profile of β-catenin has historically been challenging. When using chromatin immunoprecipitation followed by sequencing (ChIP-seq; Furey, 2012), the addition of double-crosslinking steps aimed at preserving not only the DNA-protein - but also protein-protein interactions - yielded clear β-catenin profiles both *in vitro* (Schuijers et al., 2014) and in *vivo* (Cantù et al., 2018). These protocols, however, present several limitations, including the fact that they are at risk of generating numerous artifacts and, most importantly, require enormous amounts of cells, typically in the range of tens of millions (Doumpas et al., 2019; Zimmerli et al., 2020).

Among the several technologies recently developed to profile DNA-binding proteins, Cleavage Under Targets and Release Using Nuclease (CUT&RUN, hereafter C&R) has emerged as method of choice, as it does not require crosslinking and yields genome-wide TFs profiling from a significantly lower cell input than ChIP-seq (Skene and Henikoff, 2017; Skene et al., 2018). C&R relies on the antibody-mediated recognition of specific target proteins by the fusion of proteinA/G with micrococcal nuclease (pAG-MN). pAG-MN, upon Ca^2+^ administration, cleaves the underlying DNA and generates short fragments that can be harvested, sequenced, and mapped onto a reference genome, producing TF-specific genome-wide binding patterns (Meers et al., 2019a).

However, in our attempts to characterize the genomic positioning of the WNT/β-catenin regulators, C&R systematically failed in producing a binding profile of the non-DNA-binder β-catenin. To solve this limitation, we tested a number of biochemical adaptations in select steps of the C&R protocol, to finally generate a modified procedure referred to as C&R-LoV-U (Low Volume and Urea). C&R-LoV-U utilizes nuclear extraction and *in situ* protein denaturation to improve retrieval of DNA fragments that are associated to transcriptional complexes. Reduced volumes and a streamlined pipeline further confer high reproducibility and scalability to the protocol. Importantly, C&R-LoV-U not only allows detection and profiling of β-catenin, but also all the other classes of chromatin binding proteins tested: i.e., histone modifications, classical TFs, and other components of large multi-protein complexes. We employed C&R-LoV-U on cultured HEK293T cells, as well as *ex vivo* in cells isolated from mouse developing hindlimbs which present a relatively small subset of cells with active WNT signaling (Maretto et al., 2003). We propose that C&R-LoV-U will permit the study of a broad spectrum of chromatin regulators which differ based on DNA-binding affinity, proximity to DNA, and position within large multi-protein complexes. Moreover, we envision that systematic application of this protocol will allow the study of the full set of non-DNA-binding transcriptional regulators that act downstream of onto-genetically fundamental signaling cascades.

## Results and Discussion

### *In situ* urea administration on isolated nuclei allows release of β-catenin-associated DNA fragments

CHIR stimulation in HEK293T cells generates a reproducible genome-wide DNA binding profile of β-catenin, detectable via ChIP-seq (Doumpas et al., 2019). In our hands, C&R repeatedly failed in recapitulating the β-catenin ChIP-seq profile in this cell line, generating at best sub-optimal enrichment regions that do not allow reproducible peak calling (Figure 1A). We reasoned that β-catenin might constitute a difficult C&R target mainly for three reasons. Firstly, when WNT signaling is active, β-catenin must build up in the cytosol before being translocated to the nucleus. We suspect that the high level of cytosolic β-catenin could sequester the added antibody and subsequent pAG-MN, hindering its ability to reach nuclear β-catenin and to cleave DNA. The second could be its physical distance from DNA, which might not allow pAG-MN to simultaneously reach its target (that is, the antibody against β-catenin) while cleaving the underlying genomic region. A third explanation might reside in the positioning of β-catenin within a multiprotein complex, which would hinder or retard the release into solution of any cleaved DNA fragments. We consider the second explanation unlikely, as the utilization of a secondary antibody, as previously done to extend the reach of pAG-MN (Hainer and Fazzio, 2019), did not improve the experimental outcome when tested. On the other hand, nuclear extraction combined with the addition of chaotropic agents in the elution buffer enabled the release and harvest of a considerable amount of DNA fragments that correctly mapped to WNT Responsive Elements (WRE) in the vicinity of WNT target genes (Figure 1A). Based on this finding, we developed an elution buffer in which a high final concentration of urea (up to 8.8M) proved optimal in maximizing the recovery of DNA fragments associated to the pAG-MN-dependent cleavage pattern produced when specifically targeting β-catenin (Figure 1A). Key comparisons support the reliability of our protocol modification: first is the recapitulation of classical β-catenin binding targets as observed in previous ChIP-seq assays (Figure 1B; Doumpas *et al*, 2019). Second is the reproducibility of the protocol, providing comparable signal tracks (Figure 1B) and peak enrichment across three replicates (Figure 1C) in primarily intergenic and intronic regions, as found previously (Figure 1C; Hoverter *et al*, 2014; Cantù *et al*, 2018). Third is the statistical enrichment for TCF/LEF motifs as primary transcription factor signature in the sequence underlying the high confidence β-catenin peaks (called in at least 2 of 3 replicates as described in Yang *et al*, 2014; Figure 1D). Last is the prevalence of WNT pathway-related Gene Ontology categories across peak associated gene sets (Figure 1E).

**Figure 1:**
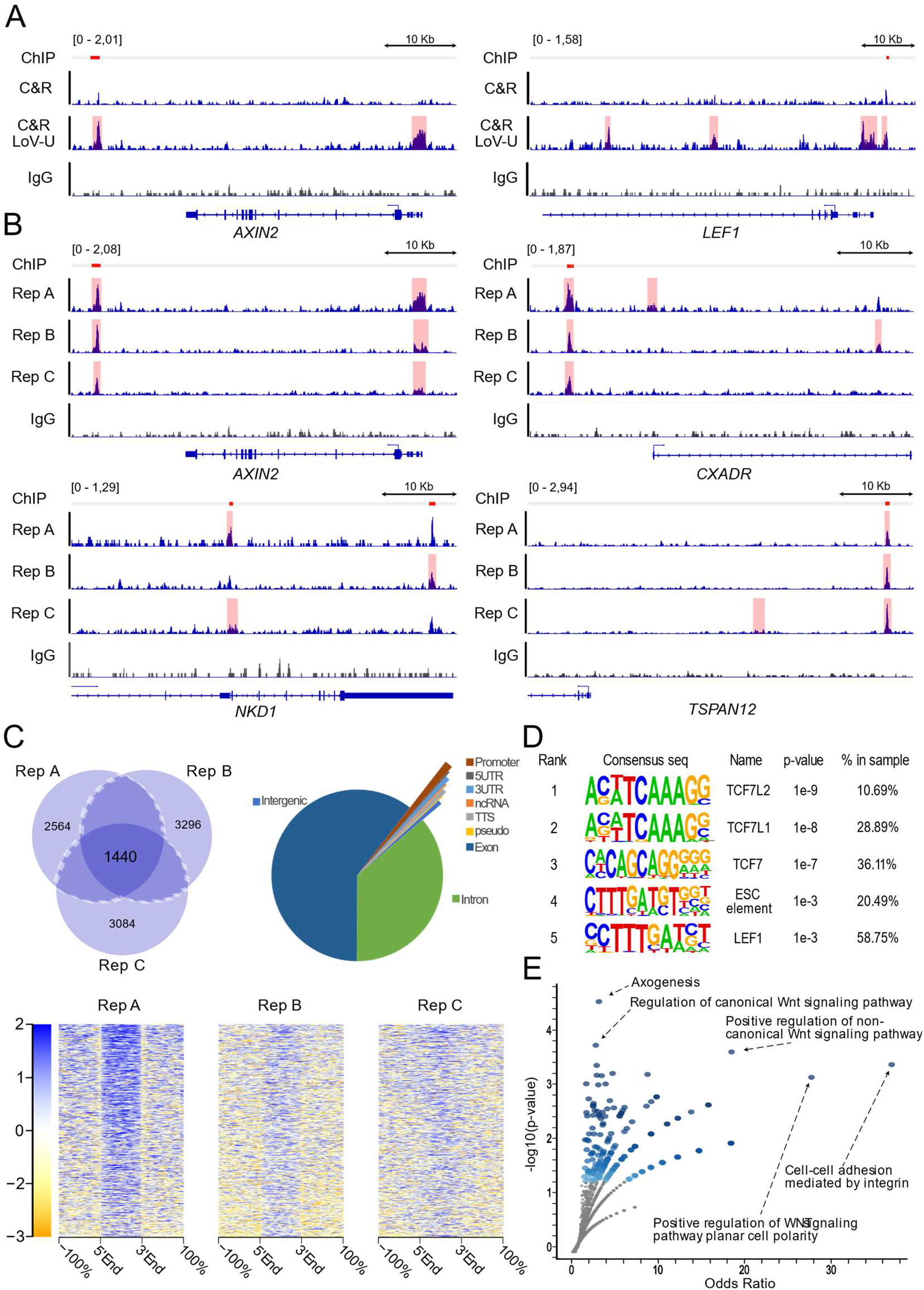
CUT&RUN with nuclear extraction and Urea-mediated release allows reproducible profiling of β-catenin binding in HEK293T. **A.** Genome coverage tracks of traditional CUT&RUN (C&R) for β-catenin and C&R using urea in the elution buffer (C&R-LoV-U: Low Volume-Urea, see Figure 2 for a complete explanation of the protocol) for β-catenin and IgG negative control, scaled to signal per million reads. Peak regions called by SEACR are shaded. C&R-LoV-U shows enriched signal compared to traditional C&R for β-catenin, and successfully recapitulates previously published ChIP-seq peaks at known WNT responsive elements, indicated by red lines corresponding to the exact positions of the β-catenin peaks called in Doumpas et al., 2019. **B.** Genome coverage tracks of 3 biological replicates of the C&R-LoV-U β-catenin and IgG negative control, showing reproducible signal enrichment across replicates. **C.** Left: Venn diagram of number of peaks called by SEACR for β-catenin replicates. The peaks called in 2 of 3 replicates were considered high-confidence and used for downstream analyses. Right: Pie chart showing genomic region annotations for β-catenin peaks. Bottom: Signal enrichment plot displaying fold-change over IgG control for each replicate over the high-confidence peak regions. **D.** Motif analysis results for high-confidence β-catenin peak regions, showing significant enrichment for TCF/LEF binding motifs. **E.** Gene Ontology analysis of the peak-associated genes, where the odds ratio (ratio of input list to reference list, x-axis) and the statistical significance (y-axis) for groups of “GO-biological processes” are represented, shows enrichment for several WNT pathway-related mechanisms.

### Reduced volumes and design of 3D-printable magnetic racks improve scalability – the Low Volume and Urea (LoV-U) protocol

During the optimization of the protocol, we strived to include additional modifications that would allow us to simultaneously streamline the procedure and improve scalability and throughput of the technique, thereby permitting the concurrent profiling of a higher number of samples (Figure 2). These modifications were mostly designed to decrease reaction volumes; for instance, reducing wash volume but increasing number of washing steps yielded similar signal-to-noise ratios, but permitted us to perform the entire protocol using 200 μl PCR tubes and multichannel pipettes. The Low Volumes (LoV) parallel processing improved the procedure speed, its cost-effectiveness (that is, it is possible to reduce the amount of reagents used proportionally to the final volume) and promoted reproducibility across replicates (Figure 2). To facilitate washes and buffer changes, CUT&RUN uses magnetic beads as a substrate and thus requires the use of a magnetic tube rack. To this purpose, we designed and 3D printed magnetic racks that provide with a better fit for the processing of 8 samples simultaneously at virtually no additional cost. As the use of these is integral part of our procedure, we have included the design of magnetic racks and all the necessary instructions for their 3Dprinting in Supplementary File 1.

**Figure 2:**
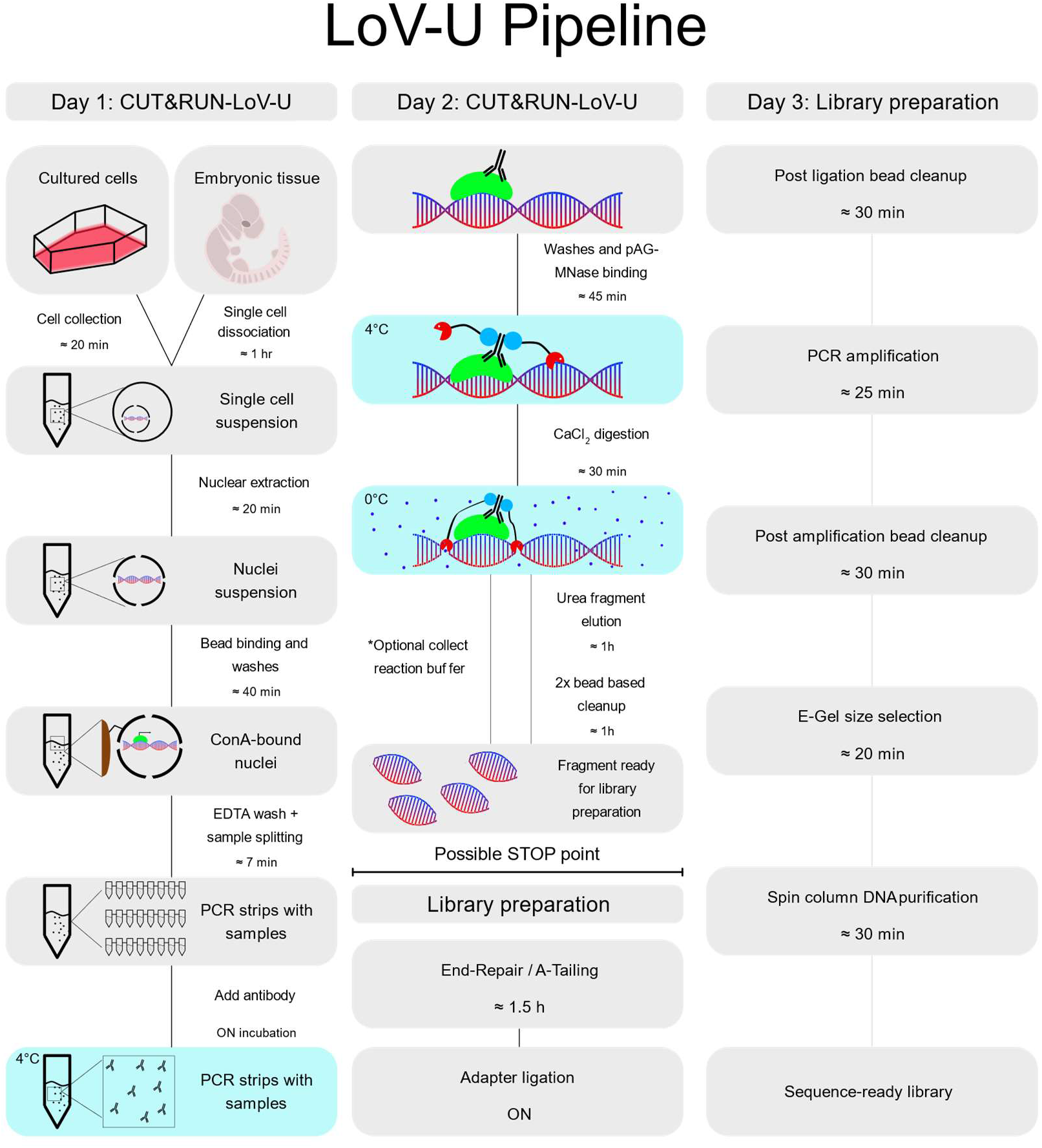
Workflow for the CUT&RUN Low Volume and Urea (C&R LoV-U) protocol. **Day 1:** cells are harvested from adherent culture or embryonic tissue and dissociated to a single cell suspension. Nuclei are extracted and bound to Concanavalin A (ConA) beads, and the sample is split into PCR tubes for each condition. Antibodies are added and samples are incubated overnight. **Day 2:** All steps are performed in PCR tubes with a multi-channel pipette. First, excess antibodies are washed away, and the samples are incubated with pAG-MN. Excess pAG-MN is removed, then calcium is added to induce pAG-MN digestion which proceeds for 30 min. A buffer containing urea is added as a chaotropic agent to the nuclei preparation, allowing *in situ* protein solubilization and release of protein-bound chromatin fragments. For factors with smaller expected fragment profiles, the digestion buffer can be retained and added back before purification. Bead-based DNA purification removes contaminants, and renders samples suitable for library preparation. End-repair and A-tailing is performed, followed by overnight adapter ligation. **Day 3:** Library preparation continues in PCR tubes. Post-ligation bead cleanup, library amplification, post-amplification bead cleanup, gel-based size selection and DNA purification can be performed in 2 – 3 hr, resulting in sequencing ready DNA libraries. With the gel-based size selection, any adapter-dimer contamination can be completely removed and DNA fragment size estimated.

### C&R-LoV-U enables profiling across different types of targets

We hypothesize that different classes of targets present different challenges to detection using C&R and similar techniques (Figure 3A). For instance, chromatin marks and histone modifications are successfully detected with extremely low number of cells likely due to their abundance and the breadth of the chromatin region exposing their epitope (Calo and Wysocka, 2013; Meers et al., 2019a). TFs, conversely, are less abundantly expressed, and are located at precise positions on the chromatin, and are more challenging to detect (Furey, 2012). Finally, non-DNA binders would constitute the most difficult type of target, as exemplified by our C&R attempts and the previous need of ChIP-seq protocols to adopt a dual cross-linking approach (Cantù et al., 2018; Salazar et al., 2019; Schuijers et al., 2014). This would suggest that our protocol could perform well also when applied to “easier” targets than β-catenin. However, we noticed that, when targeting β-catenin, C&R-LoV-U results in an enrichment of supra-nucleosomal DNA fragments (>150 nucleotides), larger than those obtained with the original C&R targeting β-catenin (Figure 2B). Classical TFs, on the other hand, typically possess a footprint of smaller, sub-nucleosomal, DNA fragments (Meers et al., 2019b). Therefore, we aimed at testing whether C&R-LoV-U could be employed for other types of targets in addition to β-catenin. As shown in Figure 3, our protocol successfully profiled the other two classes of targets: i) histone post-translational modifications (H3K4me and H3K4me2), and ii) DNA-binding TFs (the WNT signalling relevant LEF1, TCF7, TCF7L1 and TCF7L2). C&R LoV-U for H3K4me and H3K4me2 yielded the expected fragment lengths corresponding to nucleosomal and di-nucleosomal sizes, as described in Meers *et al*, 2019b, together with widespread enrichment downstream of transcriptional start sites, as described in Soares *et al*, 2017 (Figure 3C). When targeting TFs, the length distribution of the obtained fragments was consistent with average smaller size and included both sub-nucleosomal and nucleosomal fragments (Figure 3D). Moreover, when the reads were mapped to a reference genome, the binding profiles for the different TCF/LEF transcription factors partially overlapped with each other (Figure 3D), consistent with their known partial redundancy (Cadigan and Waterman, 2012; Moreira et al., 2017). Additionally, considerable overlap between the high confidence β-catenin peaks and the TCF/LEF peaks lends credibility to these datasets, while the presence of β-catenin peaks that are seemingly independent of TCF/LEF could be due to peak calling discrepancies, but aligns with previous findings (Doumpas et al., 2019) (Figure 3D). Unique peaks for the different TCF/LEF TFs are likely due to a combination of locus specificity and imperfect peak calling (Figure 3E). As shown above, our improved C&R-LoV-U protocol consistently performs on all the types of targets tested and is therefore suitable for genome-wide profiling of all chromatin-interacting proteins.

**Figure 3:**
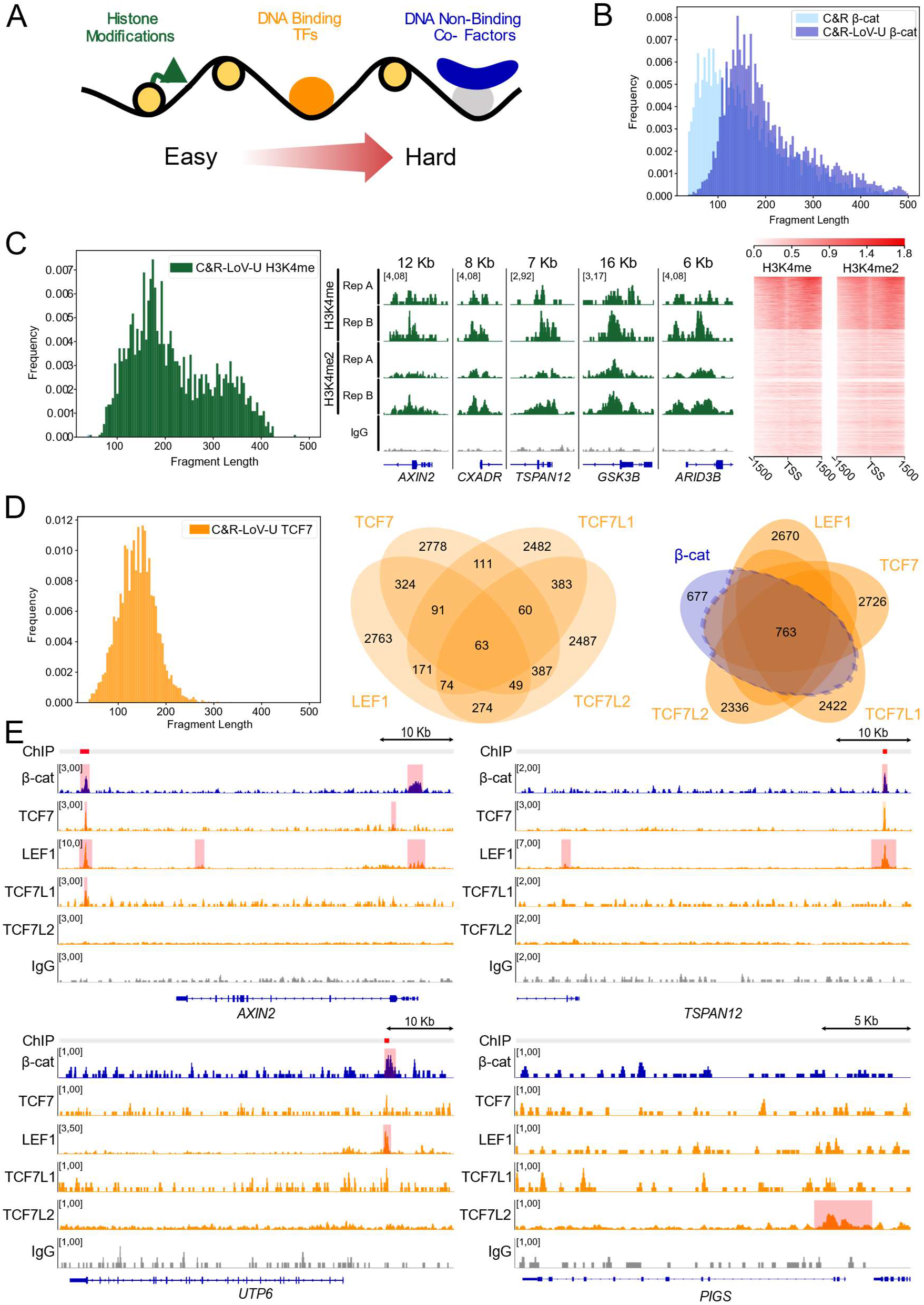
C&R-LoV-U is well adapted to the entire spectrum of chromatin associated targets. **A.** Schematic representation of the pro-posed difficulty for targets of chromatin profiling techniques. **B.** Histogram showing fragment length distribution for traditional C&R and C&R-LoV-U against β-catenin, C&R-LoV-U resulting in comparatively larger fragment size. **B.** Left: Histogram of fragment length for C&R LoV-U against H3K4me2, showing both nucleosomal (~150 bp) and di-nucleosomal (~300 bp) fragments. Middle: Genome coverage tracks scaled to signal per million reads showing 2 replicates of C&R-LoV-U for H3K4me and H3K4me2 near the transcriptional start site (TSS) of several loci. Right: Signal enrichment plots for H3K4me and H3K4me2 around the TSS for all Refseq genes. **D.** Left: Histogram of fragment length for C&R-LoV-U against TCF7, showing both subnucleosomal (<120) and nucleosomal fragments. Middle: Venn diagram of overlap of SEACR called peaks for the TCF/LEF transcription factors (n = 1). Right: Venn diagram of the overlap between high-confidence β-catenin peaks and the TCF/LEFs. Over half of β-catenin peaks are called in at least one TCF/LEF dataset. **E.** Genome coverage tracks scaled to signal per million reads showing C&R-LoV-U against β-catenin, LEF1, TCF7, TCF7L1, TCF7L2 and IgG negative control. At the *AXIN2* enhancer, peaks are called in 3 of 4 TCF/LEFs and some signal enrichment can be seen in all tracks, while other loci show preferences for one or more of the TCF/LEF factors.

### β-catenin profiling *in vivo* in developing mouse tissue uncovers tissue-specific targets

We aimed at testing the suitability of our protocol in a onto-genically relevant tissue extracted during mouse organogenesis. We selected developing hindlimbs at 11.5 days post coitum (dpc), as β-catenin is required to initiate the gene expression program toward formation of this organ (Kawakami et al., 2011), and because hindlimbs display a relatively thin layer of cells with active WNT signalling (Maretto et al., 2003). By employing C&R-Lov-U, we successfully identified 171 β-catenin peaks, present in two biological replicates, which were annotated by GREAT (McLean et al., 2010) to 179 genes. To our surprise, of these, only 12 were in common with the annotated target genes identified in HEK293T cells, while 167 appeared to be specific to the hindlimb (Figure 4A). We noticed that the stringency of our peak calling might cause a reduced overlap between the HEK293T and the hindlimb peaks, thereby leading to overestimate the tissue-specific subsets. For example, by using the same statistical parameters, *Axin2* was not called as a gene-associate peak in the hindlimb despite the consistent signal enrichment observed at its promoter (Figure 4B). Therefore, we manually searched the list for hindlimb-specific genes to examine whether previously neglected β-catenin targets might include some that are known to be relevant for formation of this organ. Interestingly, we found notable ones such as *Ets2* and *Ezh2*, both of which have been shown to be involved in mouse limb development (Ristevski et al., 2002; Wyngaarden et al., 2011). Another novel target of interest is *Kifap3* (Figure 4B). Here, the β-catenin peak does not correspond to its exact location in HEK293T (by comparing the mouse and the human homologue genomic sequences), but it lies within a different intron of the gene. Of note however, LEF1 also associates to these genomic loci (Figure 4B), thereby increasing our confidence in the reliability of these binding events even in regions of relatively low signal. It is relevant to point out that the introns of *Kifap3* have been shown to contain a limb-specific enhancer that is activated only during mouse limb formation (Nolte et al., 2014). Our data establishes that this limb-relevant ensemble of developmental regulatory regions is likely regulated by canonical WNT/β-catenin signaling. Similarly to the HEK293T peak set, the hindlimb binding regions are primarily located in intergenic and intronic regions of the genome (Figure 4C). We performed motif analysis on the hindlimb peak set and identified, in addition to TCF/LEF motifs, enrichment of the motifs of other TFs, such as GATA6 and FOXA (Figure 4C). Notably, GATA4 and GATA6 are present in an anterior-posterior gradient in the forelimb, but it is precisely GATA6 being expressed principally in hindlimb buds, where it acts as inhibitor of *Shh* (Kozhemyakina et al., 2014). Our data therefore unearths a potential GATA6-WNT/β-catenin interplay, crucial to balance the levels of SHH in ultimately determining digit patterning. We identified several other hindlimb-only target genes that display β-catenin binding in their regulatory regions, and manually checked that these regions do not show β-catenin enrichment in HEK (Supplementary Figure 1B). Among these: *Tle1*, encoding for the Transducin-Like Enhancer Of Split/Groucho repressor and known to bind on TCF/LEF at WREs and repress transcription of WNT target genes (Figure 4D)(Sekiya and Zaret, 2007); *Igf1*, known to play a role in chondrogenesis and apoptosis in the developing limb (Supplementary Figure 1A)(Kleffens et al., 1998); *Taf4b*, a basal transcription factor downstream of TGFβ signaling and highly expressed in the limbs during development (Figure 4D)(Iwata et al., 2010); *Shox2*, a transcription factor required for proper bone and muscle development in the mouse limbs (Supplementary Figure 1A) (Vickerman et al., 2011). Taken together, our datasets indicate that C&R-LoV-U possesses the sensitivity to identify both common - as well as tissue-specific - β-catenin target genes, and that application of this protocol will allow the discovery of how a seemingly universal transcriptional cascade can regulate differential sets of target genes depending on the cellular context.

**Figure 4:**
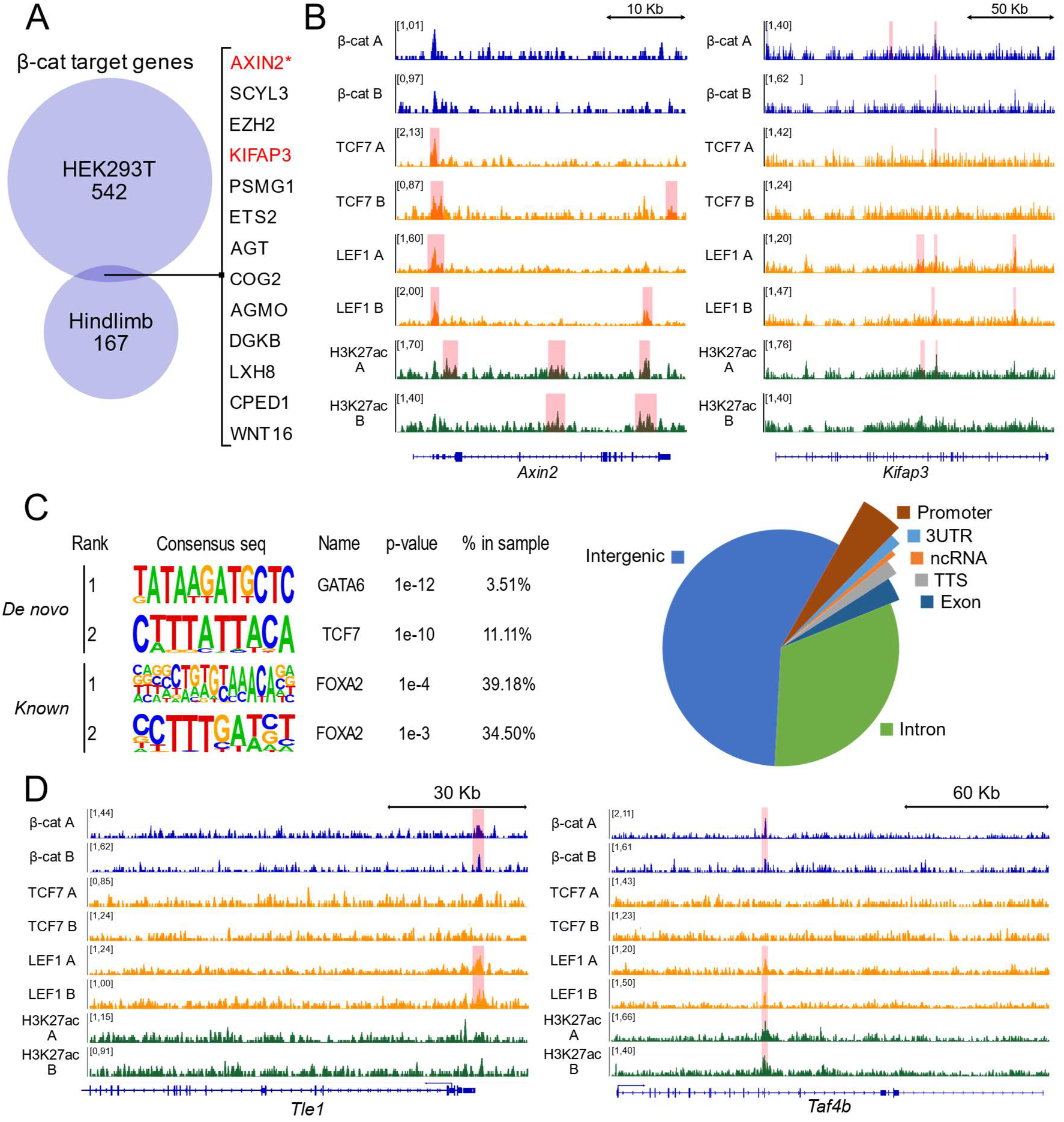
C&R-LoV-U uncovers a hindlimb specific β-catenin binding profile. **A.** Overlap of GREAT annotated target genes for HEK293T β-catenin peaks and hindlimb at 11.5 days post coitum (dpc) β-catenin peaks. Only 12 genes are shared, demonstrating high tissue-specificity of β-catenin activity. **B.** C&R-LoV-U genome coverage tracks for β-catenin, TCF7, LEF1 and H3K27ac in hindlimb at the *Axin2* and *Kifap3* loci, scaled to signal per million reads. Signal enrichment can be seen at the *Axin2* promoter and intronic enhancer regions of *Kifap3.* **C.** Left: Motif analysis of hindlimb β-catenin peaks, showing enrichment of GATA6 and FOXA factors in addition to TCF/LEF. Right: pie chart with peak region annotations of hindlimb β-catenin peaks. **D.** C&R-LoV-U genome coverage tracks for β-catenin, TCF7, LEF1 and H3K27ac in hindlimb at the *Tle1* and *Taf4b* loci, scaled to signal per million reads. Signal enrichment for β-catenin and LEF1 can be seen at the promoter of *Tle1*, while enrichment of β-catenin, LEF1 and H3K27ac can be seen within an intronic region of *Taf4b*, likely an enhancer.

## Supporting information

Supplemental Protocol 1

---

**Supplementary Figure 1:**
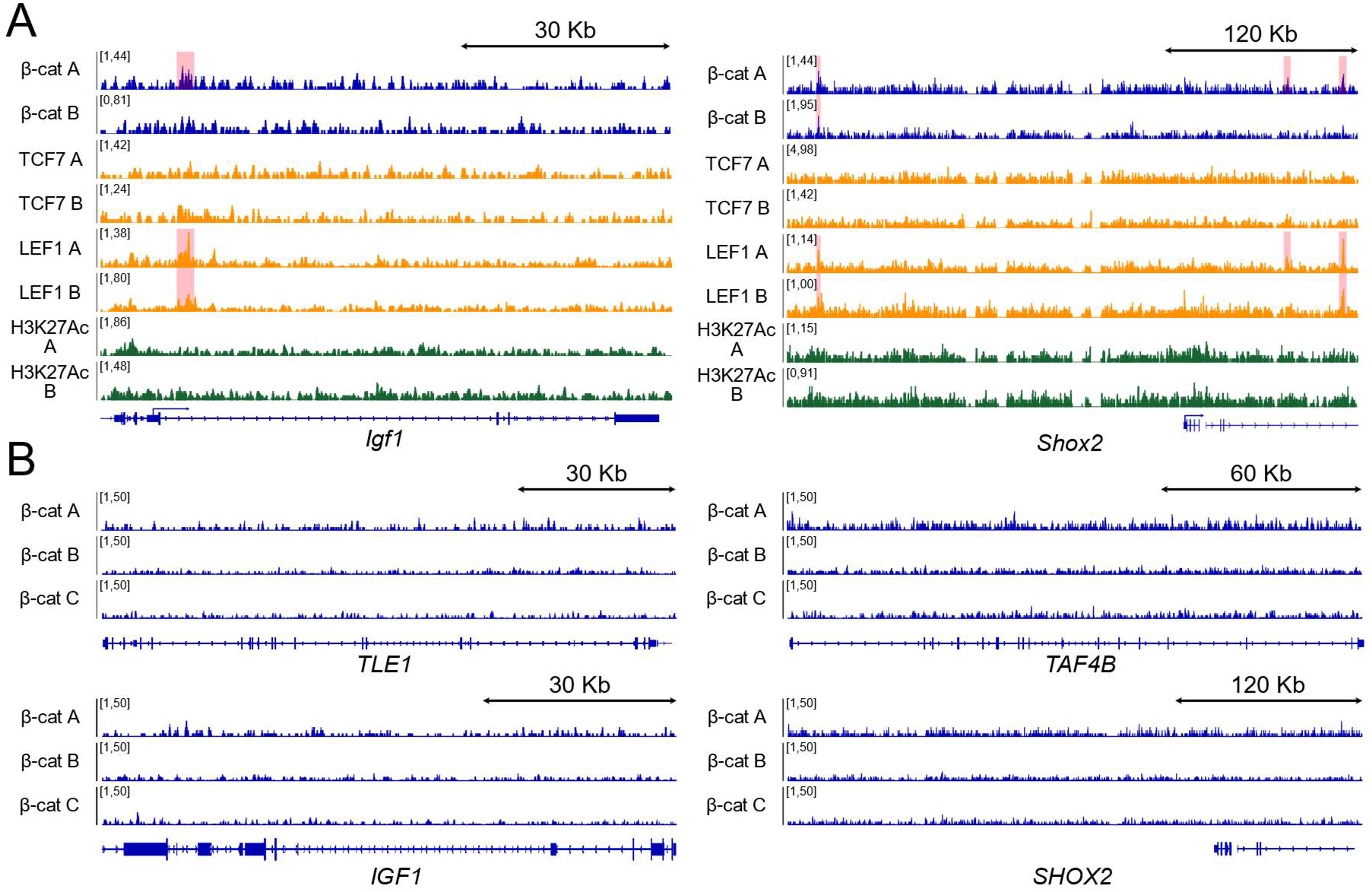
**A.** C&R-LoV-U genome coverage tracks for β-catenin, TCF7, LEF1 and H3K27ac in hindlimb at the *Igf1* and *Shox2* loci, scaled to signal per million reads. Enrichment in β-catenin and LEF1 can be seen near the promoter of *Igf1*, while the *Shox2* locus contains many sites of signal enrichment in β-catenin, LEF1 and H3K27ac, which have previously been reported as enhancers (Abassah-oppong *et al*, 2020). **B.** C&R-LoV-U genome coverage tracks for β-catenin in HEK293T, showing a lack of signal enrichment in corresponding regions to those shown in hindlimb near *Tle1, Taf4b, Igf1* and *Shox2.*

## Acknowledgements

The authors are grateful to the Henikoff Lab for donating the first batch of pA-MN protein, to the laboratories of Prof. Francisca Lottersberger and Prof. Stefan Koch for continuous scientific input, to Dr. Amaia Jauregi-Miguel for experimental help, and Dr. Stefan Pellenz for the collaborative efforts in providing antibodies suggestions. This work was supported by Grants to C.C. from Cancerfonden (CAN 2018/542 and 21 1572 Pj), and the Swedish Research Council, Vetenskapsrådet (2021-03075). C.C. is a Wallenberg Molecular Medicine (WCMM) fellow and receives generous financial support from the Knut and Alice Wallenberg Foundation. The computations and data handling were enabled by resources provided by the Swedish National Infrastructure for Computing (SNIC) at [SNIC CENTRE] partially funded by the Swedish Research Council through grant agreement no. 2018-05973.

## Author contributions

G.Z., A.N., M.J. designed and performed the experiments. P.P. assisted with the *in vivo* work and provided key improve-ments in cells preparation from mouse tissue. A.N. and M.P. carried out the bioinformatics analyses. G.Z. composed the figures. All authors assisted with data interpretation. G.Z., A.N. and C.C. wrote the manuscript. All authors contributed to multiple rounds of manuscript revision. C.C. supervised the research team and provided financial support for the study.

## Competing interest statement

The authors declare no competing interests.

## Materials and Methods

### Data availability

All data can be found at https://www.ebi.ac.uk/arrayex-press/experiments/E-MTAB-11951, accession number E-MTAB-11951.

### Cell Culture

Human embryonic kidney 293T cells (HEK293T) were cultured at 37 °C in a humidified incubator with 5% CO2. Cells were cultured in high glucose Dulbecco’s Modified Eagle Medium (Cat. #41965039, Gibco) supplemented with 10 % bovine calf serum (Cat. #1233C, Sigma-Aldrich) and 1X Penicillin-Streptomycin (Cat. #15140148, Gibco).

### Animal Experimentation and Tissue Processing

Animal housing and experimentations were performed according to the Swedish laws and guidelines under the ethical animal work license obtained by C.C. at Jordbruksverket (Dnr 2456-2019).

JAX Swiss Outbred mice (strain #: 034608) were used for all experiments. Animals were kept in Allentown NexGen IVCs, floor area 500 cm^2^. The maximum number of mice was four per cage. Cages were supplied with aspen wood shavings as bedding and two types of shredded paper as nesting material. Paper tubes were provided as additional enrichment. Temperature was set at 21 +/− 2 degrees Celsius, humidity to 45-65%, and light cycle was set to 12h/12h (7.00 am/7.00 pm). The animals had unrestricted access to sterilized drinking water, and ad libitum access to a pelleted and extruded mouse diet in the food hopper. Mice were housed in a barrier-protected specific pathogen-free unit. The specific pathogen-free status of the animals was monitored frequently and confirmed according to FELASA guidelines by a sentinel program. The mice were free of all viral, bacterial, and parasitic pathogens listed in FELASA recommendations (Mähler et al., 2014). The age of embryos was determined according to timed mating and vaginal plug observation (0.5 dpc) and confirmed by morphological criteria. Pregnant females were sacrificed by cervical dislocation and E11.5 embryos were surgically removed. Hindlimbs were dissected under a dissection stereomicroscope (SZ61, Olympus), pooled, and dissociated to single cell via incubation in Tryple Express Enzyme (12-604-013, Thermo Fisher Scientific; 1 ml /4 limbs) for 15 minutes at 37 °C on a shaker. The cell suspension was resuspended in ice-cold PBS, filtered through a 40 μm cell strainer (KKE3.1, Carl Roth) and further processed for CUT&RUN.

### CUT&RUN

CUT&RUN was performed as described in Skene *et al*, 2018. Prior to harvest, HEK293T cells were cultured in media containing 10 μM CHIR99021 (Cat. #SML1046, Sigma Aldrich) for 24 hr. 500,000 cells were harvested using TrypLE (Cat. #12604013, Gibco), washed twice in wash buffer (HEPES pH 7.5 [20 mM], NaCl [150 mM], Spermidine [0.5 mM], Roche Complete Protease Inhibitor EDTA-Free (Cat. # CO-EDTAFRO, Roche)) and bound to 20 μl Magnetic ConA Agarose beads (Cat. #ABIN6952467, antibodies-online) equilibrated in binding buffer (HEPES pH 7.5 [20 mM], KCl [10 mM], CaCl_2_ [1 mM], MnCl_2_ [1 mM]). Antibody incubation was performed in 150 μl antibody buffer (wash buffer with digitonin [0.01%] and EDTA [2 mM]) with 1.5 μl anti-β-catenin antibody (Cat. #ABIN2855042, antibodies-online) overnight at 4 °C. Thereafter washes were performed with wash buffer containing 0.01% digitonin. After overnight incubation, samples were washed three times and pAG-MN protein was added at 0.6 μg/ml, in total volume of 150 μl. pAG/MNase was a gift from Steven Henikoff (Addgene plasmid #123461; http://n2t.net/addgene:123461; RRID: Addgene_123461), expressed and purified as described in (Meers et al., 2019a). Samples were washed three times, followed by digestion for 30 min in wet ice in wash buffer supplemented with 2 mM CaCl_2_. 2X stop buffer (NaCl [340 mM], EDTA [20 mM], EGTA [4 mM], digitonin [0.05%], RNase A [100 μg/ml], glycogen [50 μl/ml]) was added to stop the reaction, and samples were incubated for 30 min at 37 °C. Beads were collected on the magnet and liquid transferred to a new tube. 2.5 μl proteinase K (Cat. #P8107S, New England BioLabs) and 2 μl 10% SDS were added, and samples incubated for 1 hr at 50 °C. 200 μl phenol-chloroform (Cat. #P3803, Sigma Aldrich) was added and samples were vortexed before being centrifuged at 15,000 rcf for 10 minutes. The aqueous phase was transferred to a new tube containing 1.5 μl glycogen, 20 μl 3M sodium acetate and 500 μl 100% ethanol. Samples were incubated ON at −20 °C. DNA precipitation was performed by centrifugation at 15,000 rcf for 45 min at 4 °C. DNA pellets were washed once in 70% ethanol and centrifuged for 10 min at 15,000 rcf. Pellets were allowed to air dry before being resuspended in 20 μl Tris-HCl pH 7.5.

### CUT&RUN LoV-U

The detailed protocol is attached as Supplementary Protocol 1. HEK293T cells were cultured in media containing 10 μM CHIR99021 for 24 hr, then 500,000 cells/sample were harvested using TrypLE and washed twice in DPBS (Cat. #14190094, Thermo Fisher Scientific). For each ex *vivo* sample ~100,000 cells (3 hindlimbs at 11.5) were collected. Cells were washed three times in Nuclear Extraction (NE) buffer (HEPES-KOH pH-8.2 [20 mM], KCl [10 mM], Spermidine [0.5 mM], IGEPAL [0.05%], Glycerol [20%], Roche Complete Protease Inhibitor EDTA-Free), resuspended in 40 μl NE per sample and bound to 20 μl Magnetic ConA Agarose beads equilibrated in binding buffer. After incubation nuclei and beads are resuspended for 5 min in EDTA wash buffer (wash buffer with EDTA [0.2 mM]). Samples were divided in 200 μl PCR tubes and antibody incubation was performed in 200 μl wash buffer with 2 μl of antibody (for antibody and batch information, see Table 1) ON at 4 °C on a rotator. After ON incubation samples were washed 5 times in wash buffer and resuspended in 200 μl of pAG-MN buffer (wash buffer with pAG-MN 0.6 μg/ml) for 30 min at 4 °C on a rotator. Samples were washed five times, followed by digestion for 30 min in wet ice in wash buffer with 2 mM CaCl_2_. After 30 min the digestion buffer was removed and the reaction was stopped with 50 μl of 1X Urea STOP buffer (NaCl [100 mM], EDTA [2 mM], EGTA [2 mM], IGEPAL [0.5%], Urea [8.8 M]) and the samples were incubated 1 hr at 4°C. Beads were collected on the magnet and liquid transferred to a new PCR tube in which it was cleaned up twice using Mag-Bind TotalPure NGS beads (Cat. #M1327, Omega Bio-Tek) at 2X, and then resuspended in 20 μl Tris-HCl pH 7.5.

**Table 1:** Antibody Information.

| Target | Species | Clonality | Supplier | Cat. # | Batch/Lot |
| --- | --- | --- | --- | --- | --- |
| β-catenin | Rabbit | poly | antibodies-online | ABIN2855042 | 42564 |
| LEF1 | Rabbit | poly | antibodies-online | ABIN1680678 | 0003180201 |
| TCF7 | Rabbit | poly | antibodies-online | ABIN5620945 | X21031510 |
| TCF7L1 | Rabbit | poly | antibodies-online | ABIN6972849 | 20011001 |
| TCF7L2 | Rabbit | mono | antibodies-online | ABIN6945139 | BA04368866 |
| H3K4me | Rabbit | poly | antibodies-online | ABIN3023251 | 3560036504 |
| H3K4me2 | Rabbit | poly | Invitrogen | 710796 | 2059496 |
| H3K27Ac | Rabbit | poly | antibodies-online | ABIN6971893 | 06921014 |
| Rabbit IgG | Guinea | poly | antibodies-online | ABIN101961 | NE-200-10200 |

### Library Preparation and Sequencing

Library preparation was performed using the KAPA Hyper Prep Kit for Illumina platforms (Cat. #KK8504, KAPA Biosystems) according to the manufacturer’s guidelines with the following modifications. End repair and A-tailing was performed with 0.4X volume reactions with 20 μl of purified DNA. The thermocycler conditions were set to 12 °C for 15 min, 37 °C for 15 min and 58 °C for 25 min to prevent thermal degradation of the shortest fragments. Adapter ligation was performed in 0.4X volume reactions. KAPA Dual Indexed adapters were used at 0.15 μM. A post-ligation cleanup was performed with Mag-Bind TotalPure NGS beads at 1.2X the sample volume. Resuspension was done in 10 mM Tris-HCl pH 8.0. Library amplification was performed in 0.5X volume reactions. The cycling conditions were set as follows: 45 sec initial denaturation at 98 °C, 15 sec denaturation at 98 °C, 10 sec annealing/elongation at 60 °C, 1 min final extension at 72 °C, hold at 4 °C, with 13 cycles. After amplification, a post-amplification cleanup was performed using NGS beads at 1.2X sample volume. Libraries were then run on an E-Gel EX 2% agarose gel (Cat. # G402022, Invitrogen) for 10 min using the E-Gel Power Snap Electrophoresis System (Invitrogen). Bands of interest between 150 and 500 bp were excised and purified using the GeneJET Gel Extraction Kit (Cat. #K0691, Thermo Scientific) according to manufacturer’s instructions. Libraries were quantified with the Qubit (Thermo Scientific) using the high sensitivity DNA kit (Cat #Q32854, Thermo Scientific), pooled and sequenced 36 bp pair-end on the NextSeq 550 (Illumina) using the Illumina NextSeq 500/550 High Output Kit v2.5 (75 cycles) (Cat. #20024906, Illumina) to an approximate depth of 5 – 10 million reads per sample.

### Data Analysis

Quality of reads was assessed using fastqc (Brandine and Smith, 2022, version 0.11.9). Trimming was performed using bbmap bbduk (Bushnell *et al*, 2017, version 38.18) removing adapters, artifacts, poly AT and TA repeats, and poly G and C repeats. Reads were aligned to the hg38 genome or mm10 genome using bowtie (Langmead *et al*, 2009, version 1.0.0) with the options -v 0 -m 1 -X 500. Samtools (Li et al., 2009), version 1.11) view, fixmate, markdup and sort were used to create bam files, mark and remove duplicates, and sort bam files. Fragment size analysis was done using deeptools (Ramírez *et al*, 2016, version 3.5.1-0) bam-PEFragmentsize-hist. Bedgraphs were created using bedtools (Quinlan & Hall, 2010, version 2.23.0) genomecov on pair-end mode. Normalized signal per million reads tracks for visualization were created by removing mitochondrial reads from bam files with awk followed by using the --SPRM function of macs2 (Zhang *et al*, 2008, version 2.2.6) with the options -f BAMPE --keep-dup all --SPMR and -bdg. Peaks were called using SEACR (Meers *et al*, 2019c, version 1.3) for each bedgraph using the settings norm and stringent with a threshold set to 0.001. Final peak sets were generated by using bedtools subtract with the option -A to remove blacklisted regions (Amemiya et al., 2019), and regions overlapping with peaks called in the corresponding IgG negative control for HEK293T samples. Venn diagrams and overlap peak sets were created using Intervene (Khan & Mathelier, 2017, version 0.6.4). High confidence β-catenin peaks were considered those called in at least 2 of 3 replicates in HEK293T. For comparison with ChIP, high-confidence peak regions were downloaded from Doumpas *et al*, 2019 and converted from hg19 to hg38 using the UCSC Lift-Over (Hinrichs et al., 2006) with default settings. Signal intensity plots for β-catenin were created using ngsplot (Shen *et al*, 2014, version 2.63) with options -G hg38 -R bed -N 2 - SC global -IN 0 -CD 1 for each β-catenin replicate against the IgG negative control, and for H3K4me and H3K4me2 with the options -G hg38 -R tss -SC global -L 1500. Motif analysis was done using Homer (Heinz *et al*, 2010, version 4.11) findMotifsGenome to find motifs in the hg38 or mm10 genome. Peak set gene annotation was done using GREAT (McLean *et al*, 2010, version 4.0.4) with default parameters. The Enrichr web server (Kuleshov et al., 2016) was used for gene ontology, the Appyter option used to create the volcano plot for enriched GO biological processes using default settings.

## Notes

https://www.ebi.ac.uk/arrayexpress/experiments/E-MTAB-11951

### Competing Interest Statement

The authors have declared no competing interest.

## References

Amemiya, H.M., Kundaje, A., and Boyle, A.P. (2019). The ENCODE Blacklist: Identification of Problematic Regions of the Genome. Sci. Rep. 9, 1–5.

Brandine, G.D.S., and Smith, A.D. (2022). Falco: high-speed FastQC emulation for quality control of sequencing data [version 2; peer review: 2 approved].

Bushnell, B., Rood, J., and Singer, E. (2017). BBMerge – Accurate paired shotgun read merging via overlap. 1–15.

Cadigan, K.M., and Waterman, M.L. (2012). TCF/LEFs and Wnt Signaling in the Nucleus. Cold Spring Harb. Perspect. Biols. 4, a007906–a007906.

Calo, E., and Wysocka, J. (2013). Modification of Enhancer Chromatin: What, How, and Why? Mol. Cell 49, 825–837.

Cantù, C., Felker, A., Zimmerli, D., Prummel, K.D., Cabello, E.M., Chiavacci, E., Méndez-Acevedo, K.M., Kirchgeorg, L., Burger, S., Ripoll, J., et al. (2018). Mutations in Bcl9 and Pygo genes cause congenital heart defects by tissue-specific perturbation of Wnt/β-catenin signaling. Genes Dev. 32, 1443–1458.

Doumpas, N., Lampart, F., Robinson, M.D., Lentini, A., Nestor, C.E., Cantù, C., and Basler, K. (2019). TCF / LEF dependent and independent transcriptional regulation of Wnt/β-catenin target genes. EMBO J. 38, e98873.

Furey, T.S. (2012). ChIP-seq and beyond: New and improved methodologies to detect and characterize protein-DNA interactions. Nat. Rev. Genet. 13, 840–852.

Hainer, S.J., and Fazzio, T.G. (2019). High resolution chromatin profiling using CUT&RUN. Curr Protoc Mol Biol.

Heinz, S., Benner, C., Spann, N., Bertolino, E., Lin, Y.C., Laslo, P., Cheng, J.X., Murre, C., Singh, H., and Glass, C.K. (2010). Simple Combinations of Lineage-Determining Transcription Factors Prime cis -Regulatory Elements Required for Macrophage and B Cell Identities. Mol. Cell 38, 576–589.

Hinrichs, A.S., Karolchik, D., Baertsch, R., Barber, G.P., Bejerano, G., Clawson, H., Diekhans, M., Furey, T.S., Harte, R.A., Hsu, F., et al. (2006). The UCSC Genome Browser Database: update 2006. 34, 590–598.

Hoverter, N.P., Zeller, M.D., McQuade, M.M., Garibaldi, A., Busch, A., Selwan, E.M., Hertel, K.J., Baldi, P., and Waterman, M.L. (2014). The TCF C-clamp DNA binding domain expands the Wnt transcriptome via alternative target recognition. Nucleic Acids Res. 42, 13615–13632.

Huber, A.H., Nelson, W.J., and Weis, W.I. (1997). Threedimensional structure of the armadillo repeat region of β-catenin. Cell 90, 871–882.

Iwata, J., Hosokawa, R., Sanchez-lara, P.A., Urata, M., Slavkin, H., and Chai, Y. (2010). Transforming Growth Factor-b Regulates Basal Transcriptional Regulatory Machinery to Control Cell Proliferation and Differentiation in Cranial Neural. J. Biol. Chem. 285, 4975–4982.

Kawakami, Y., Marti, M., Kawakami, H., Itou, J., Quach, T., Johnson, A., Sahara, S., O’Leary, D.D.M., Nakagawa, Y., Lewandoski, M., et al. (2011). Islet1-mediated activation of the β-catenin pathway is necessary for hindlimb initiation in mice. Development 138, 4465–4473.

Khan, A., and Mathelier, A. (2017). Intervene: A tool for intersection and visualization of multiple gene or genomic region sets. BMC Bioinformatics 18, 1–8.

Kleffens, M. Van, Groffen, C., Rosato, R.R., and Eijnde, S.M. Van Den (1998). mRNA expression patterns of the IGF system during mouse limb bud development, determined by whole mount in situ hybridization. 138, 151–161.

Klemm, S.L., Shipony, Z., and Greenleaf, W.J. (2019). Chromatin accessibility and the regulatory epigenome. Nat. Rev. Genet. 20, 207–220.

Kozhemyakina, E., Ionescu, A., and Lassar, A.B. (2014). GATA6 Is a Crucial Regulator of Shh in the Limb Bud. 10.

Kramps, T., Peter, O., Brunner, E., Nellen, D., Froesch, B., Chatterjee, S., Murone, M., Züllig, S., and Basler, K. (2002). Wnt/wingless signaling requires BCL9/legless-mediated recruitment of pygopus to the nuclear beta-catenin-TCF complex. Cell 109, 47–60.

Kuleshov, M. V, Jones, M.R., Rouillard, A.D., Fernandez, N.F., Duan, Q., Wang, Z., Koplev, S., Jenkins, S.L., Jagodnik, K.M., Lachmann, A., et al. (2016). Enrichr: a comprehensive gene set enrichment analysis web server 2016 update. 44, 90–97.

Langmead, B., Trapnell, C., Pop, M., and Salzberg, S.L. (2009). Ultrafast and memory-efficient alignment of short DNA sequences to the human genome. 10.

Li, H., Handsaker, B., Wysoker, A., Fennell, T., Ruan, J., Homer, N., Marth, G., Abecasis, G., and Durbin, R. (2009). The Sequence Alignment/Map format and SAMtools. Bioinformatics 25, 2078–2079.

Mähler, M., Berar, M., Feinstein, R., Gallagher, A., Illgen-Wilcke, B., Pritchett-Corning, K., and Raspa, M. (2014). FELASA recommendations for the health monitoring of mouse, rat, hamster, guinea pig and rabbit colonies in breeding and experimental units. Lab. Anim. 48, 178–192.

Maretto, S., Cordenonsi, M., Dupont, S., Braghetta, P., Broccoli, V., Hassan, A.B., Volpin, D., Bressan, G.M., and Piccolo, S. (2003). Mapping Wnt/beta-catenin signaling during mouse development and in colorectal tumors. 100, 4–9.

McLean, C.Y., Bristor, D., Hiller, M., Clarke, S.L., Schaar, B.T., Lowe, C.B., Wenger, A.M., and Bejerano, G. (2010). GREAT improves functional interpretation of cis-regulatory regions. Nat. Biotechnol. 28, 495–501.

Meers, M.P., Bryson, T.D., Henikoff, J.G., and Henikoff, S. (2019a). Improved CUT&RUN chromatin profiling tools. Elife 8, 1–16.

Meers, M.P., Janssens, D.H., and Henikoff, S. (2019b). Pioneer Factor-Nucleosome Binding Events during Differentiation Are Motif Encoded. Mol. Cell 75, 562–575.e5.

Meers, M.P., Tenenbaum, D., and Henikoff, S. (2019c). Peak calling by Sparse Enrichment Analysis for CUT&RUN chromatin profiling. Epigenetics and Chromatin 12, 1–11.

Merika, M., and Thanos, D. (2001). Enhanceosomes. Curr. Opin. Genet. Dev. 11, 205–208.

Mitsis, T., Efthimiadou, A., Bacopoulou, F., Vlachakis, D., Chrousos, G.P., and Eliopoulos, E. (2020). Transcription factors and evolution: An integral part of gene expression (Review). World Acad. Sci. J. 2, 3–8.

Moreira, S., Polena, E., Gordon, V., Abdulla, S., Mahendram, S., Cao, J., Blais, A., Wood, G.A., Dvorkin-Gheva, A., and Doble, B.W. (2017). A Single TCF Transcription Factor, Regardless of Its Activation Capacity, Is Sufficient for Effective Trilineage Differentiation of ESCs. Cell Rep. 20, 2424–2438.

Mosimann, C., Hausmann, G., and Basler, K. (2009). Beta-catenin hits chromatin: regulation of Wnt target gene activation. Nat. Rev. Mol. Cell Biol. 10, 276–286.

Nikolov, D.B., and Burley, S.K. (1997). RNA polymerase II transcription initiation: A structural view. Proc. Natl. Acad. Sci. U. S. A. 94, 15–22.

Nolte, M.J., Wang, Y., Deng, J.M., Swinton, P.G., Wei, C., Guindani, M., Schwartz, R.J., and Behringer, R.R. (2014). Functional analysis of limb transcriptional enhancers in the mouse. Evol. Dev. 16, 207–223.

Quinlan, A.R., and Hall, I.M. (2010). BEDTools: A flexible suite of utilities for comparing genomic features. Bioinformatics 26, 841–842.

Ramírez, F., Ryan, D.P., Grüning, B., Bhardwaj, V., Kilpert, F., Richter, A.S., Heyne, S., Dündar, F., and Manke, T. (2016). deepTools2: a next generation web server for deepsequencing data analysis. Nucleic Acids Res. 44, W160–W165.

Rim, E.Y., Clevers, H., and Nusse, R. (2022). The Wnt Pathway: From Signaling Mechanisms to Synthetic Modulators. Annu. Rev. Biochem. 91, 1–28.

Ristevski, S., Tam, P.P.L., Hertzog, P.J., and Kola, I. (2002). Ets2 is expressed during morphogenesis of the somite and limb in the mouse embryo. 116, 165–168.

Salazar, V.S., Capelo, L.P., Cantù, C., Zimmerli, D., Gamer, L., Ohte, S., Economides, A., Carey, D.J., Feigenson, M., Pregizer, S., et al. (2019). Reactivation of a developmental Bmp2 signaling center is required for therapeutic control of the periosteal niche in the murine skeleton. Elife 8, 1–31.

Schuijers, J., Mokry, M., Hatzis, P., Cuppen, E., and Clevers, H. (2014). Wnt-induced transcriptional activation is exclusively mediated by TCF/LEF. EMBO J. 33, 146–156.

Sekiya, T., and Zaret, K.S. (2007). Repression by Groucho/TLE/Grg Proteins: Genomic Site Recruitment Generates Compacted Chromatin In Vitro and Impairs Activator Binding In Vivo. Mol. Cell 28, 291–303.

Shen, L., Shao, N., Liu, X., and Nestler, E. (2014). ngs. plot: Quick mining and visualization of next-generation sequencing data by integrating genomic databases. 1–14.

Skene, P.J., and Henikoff, S. (2017). An efficient targeted nuclease strategy for high-resolution mapping of DNA binding sites. Elife 6.

Skene, P.J., Henikoff, J.G., and Henikoff, S. (2018). Targeted in situ genome-wide profiling with high efficiency for low cell numbers. Nat. Protoc. 13, 1006–1019.

Soares, L.M., He, P.C., Chun, Y., Suh, H., Kim, T.S., and Buratowski, S. (2017). Determinants of Histone H3K4 Methylation Patterns. Mol. Cell 68, 773–785.e6.

Söderholm, S., and Cantù, C. (2020). The WNT/β-catenin dependent transcription: A tissue-specific business. WIREs Syst. Biol. Med. 1–41.

Takemaru, K.-I., and Moon, R.T. (2000). The transcriptional coactivator Cbp interacts with beta-catenin to activate gene expression. J. Cell Biol. 149, 249–254.

van Tienen, L.M., Mieszczanek, J., Fiedler, M., Rutherford, T.J., Bienz, M., Labhart, T., Desplan, C., Hursh, D., Jones, T., Bejsovec, A., et al. (2017). Constitutive scaffolding of multiple Wnt enhanceosome components by Legless/BCL9. Elife 6, 477–488.

Valenta, T., Hausmann, G., and Basler, K. (2012). The many faces and functions of β-catenin. EMBO J. 31, 2714–2736.

Vickerman, L., Neufeld, S., and Cobb, J. (2011). Shox2 function couples neural, muscular and skeletal development in the proximal forelimb. Dev. Biol. 350, 323–336.

Wyngaarden, L.A., Delgado-olguin, P., Su, I., Bruneau, B.G., and Hopyan, S. (2011). Ezh2 regulates anteroposterior axis specification and proximodistal axis elongation in the developing limb. 3767, 3759–3767.

Yang, Y., Fear, J., Hu, J., Haecker, I., Zhou, L., Renne, R., Bloom, D., and Mcintyre, L.M. (2014). Leveraging biological replicates to improve analysis in ChIP-seq experiments. Comput. Struct. Biotechnol. J.

Zhang, Y., Liu, T., Meyer, C.A., Eeckhoute, J., Johnson, D.S., Bernstein, B.E., Nussbaum, C., Myers, R.M., Brown, M., Li, W., et al. (2008). Model-based analysis of ChIP-Seq (MACS). Genome Biol. 9.

Zimmerli, D., Borrelli, C., Jauregi-Miguel, A., Söderholm, S., Brütsch, S., Doumpas, N., Reichmuth, J., Murphy-Seiler, F., Aguet, M., Basler, K., et al. (2020). TBX3 acts as tissuespecific component of the Wnt/β-catenin enhanceosome. Elife 1–17.

