## Supplemental Protocol 1 for "A New CUT&RUN Low Volume-Urea (LoV-U) protocol uncovers Wnt/β-catenin tissue-specific genomic targets"

### CUT&RUN
### LoV-U

Cantù Lab

2022

#### Buffer composition

- Nuclear extraction buffer:
  - – Hepes-KOH pH 8.2 [20 mM]
  - – KCl [10 mM]
  - – Spermidine [0.5 mM]
  - – IGEPAL 0.05%
  - – Glycerol 20%
  - – Roche Complete Proteinase inhibitor EDTA-Free
- Wash Buffer:
  - – Hepes pH 7.5 [20 mM]
  - – NaCl [150 mM]
  - – Spermidine [0.5 mM]
  - – Roche Complete Proteinase inhibitor EDTA-Free
- EDTA wash buffer:
  - – Wash buffer
  - – EDTA [0.2 mM]
- 1x Urea STOP buffer:
  - – NaCl [100 mM]
  - – EDTA [2 mM]
  - – EGTA [2 mM]
  - – NP-40 (IGEPAL) 0.5%
  - – Urea [8.8 M]
- Binding Buffer:
  - – $MnCl_2$[1 mM]
  - – $CaCl_2$[1 mM]
  - – KCl [10 mM]
  - – HEPES pH 7.5 [20 mM]

### Buffer preparation

- Nuclear extraction buffer (100ml):
  - 2ml Hepes-KOH [1 M] pH 8.2
  - 1ml KCl [1 M]
  - 25µl Spermidine [2 M]
  - 50µl IGEPAL 100%
  - 40ml Glycerol 50%
  - 2 Roche complete Proteinase inhibitor EDTA-Free;
  - 57ml dd$H_2O$
- Wash Buffer (50ml):
  - 1ml Hepes [1 M] pH 7.5
  - 1,5ml NaCl [5 M]
  - 12,5µl Spermidine [2 M]
  - 1 Roche complete Proteinase inhibitor EDTA-Free
  - 47ml dd$H_2O$
- EDTA wash Buffer:
  - 2 ml Wash Buffer
  - 8 µl EDTA 0.5 M
- 1x Urea STOP buffer (5ml):
  - 100µl NaCl [5 M]
  - 20µl EDTA [0.5 M]
  - 20µl EGTA [0.5 M]
  - 25µl NP-40 (IGEPAL) 100%
  - 4,8ml Urea [8.8 M]
- Binding Buffer (20ml last 6month):
  - 20µl $MnCl_2$[1 M]
  - 20µl $CaCl_2$[1 M]
  - 200µl KCl [1 M]
  - 400µl HEPES [1 M] pH 7.5
  - 19,36ml dd[$H_2O$]

### 1 Nuclei isolation

1.1 Harvest the cells ($5x10^4$ - $5x10^5$ cells per sample)

1.1.1 Adherent cells:
- 1.1.1.1 Wash with warm DPBS
- 1.1.1.2 Detach with Trypsin or TrypLE™
- 1.1.1.3 Stop the reaction with PBS or media with serum
- 1.1.1.4 Pellet the cells and resuspend in PBS

1.1.2 If working with embryonic tissue (e.g. Forelimbs):
- 1.1.2.1 Dissect the embryo to collect the tissue
- 1.1.2.2 Resuspend the tissue pieces in a large volume of TrypLE™(6-10 ml)
- 1.1.2.3 Incubate 15’ at 37°C on a shaker
- 1.1.2.4 Filter the cells using a cell strainer (40 µm)
- 1.1.2.5 Pellet the cells and resuspend in PBS

1.2 Spin down the cells 5’ 800 rcf and discard the liquid

1.3 Resuspend in 2 ml of NE buffer by gentle pipetting and repeat the NE wash 2X

1.4 Spin down the cells 5’ 800 rcf

1.5 Resuspend the nuclei in ≈ 20 µl of NE buffer per sample

1.6 Move the nuclei to a 2 ml microcentrifuge tube and store in ice

### 2 Concanavalin A beads binding (ABIN6952467):

2.1 In a new 2 ml microcentrifuge tube prepare 20 µl of beads per sample and add 1 ml binding buffer

2.2 Place the beads on the magnet rack and when the liquid is clear discard the supernatant

2.3 Wash the beads 2X with 1ml of binding buffer

2.4 Place the beads on the magnet rack and when the liquid is clear discard the supernatant

2.5 Resuspend the beads in ≈ 20 µl of binding buffer for each sample

2.6 Gently add the beads to the nuclei and pipette up and down to mix them

2.7 Incubate 15’ at 4°C on a shaker

#### 3 Primary antibody binding:

3.1 Place the 2 ml microcentrifuge tube on the magnet rack and when the liquid is clear discard the supernatant

3.2 Resuspend the sample in 100 µl of wash buffer per sample and split them into 200 µl tubes (use PCR strips to facilitate the next steps)

3.3 Put the tubes on the magnet rack and when the liquid is clear discard the supernatant

3.4 Resuspend the samples in 200 µl of EDTA wash buffer and incubate for 5’ RT

3.5 Place the sample on the magnet rack and when the liquid is clear discard the supernatant

3.6 Resuspend each sample in 200 µl of wash buffer

3.7 Add to each tube the right antibody (2 µl, final concentration 1:100)

3.8 Incubate ON at 4˚C or 1h 37˚C (To enhance efficiency put the tubes on a slow rotator at 4˚C).

#### 4 Secondary antibody binding (optional):

4.1 Place PCR tubes on the magnet rack and when the liquid is clear discard the supernatant (from this point we use a multi-channel pipette);

4.2 Wash the samples 5x with 200 µl of wash buffer (pipette up and down 5X to resuspend the beads)

4.3 After the washes resuspend all the samples in 200 µl of wash buffer and add to sample 1:100 secondary antibody (2 µl)

4.4 Incubate 1h in the fridge 4˚C

#### 5 pAG-MNase binding:

5.1 Place the sample on the magnet rack and when the liquid is clear discard the supernatant;

5.2 Wash the sample 5x with 200 µl of wash buffer (pipette up and down 5X to resuspend the beads in between washes);

5.3 Prepare a 1.5 microcentrifuge tube with 200 µl of wash buffer + 0.12 µg pAG-MNase per sample (pAG-mix)

5.4 Place the PCR strip with the samples on the magnet rack and when the liquid is clear discard the supernatant

5.5 Resuspend each sample in 200 µl of pAG mix

5.6 Incubate the samples 30' at 4°C in the fridge (To enhance efficiency put the tubes on a slow rotator)

5.7 Place the sample on the magnet rack and when the liquid is clear discard the supernatant

5.8 Wash the sample 5X with 200 µl of wash buffer

5.9 Place the sample on the magnet rack and when the liquid is clear discard the supernatant

5.10 Resuspend the sample in 100 µl of wash buffer.

Note: If the target has a small expected foot print (e.g. 60 bp), resuspend the samples and perform the digestion in 50 µl.

### 6 Digestion and fragment release:

6.1 Equilibrate the sample 5' at 0°C (put them in ice)

6.2 Always in ice, add $CaCl_2$ [100 mM] 1:50 to each tube (2 µl if 100 µl, 1 µl if 50 µl final volume) and mix well:

- If you have a lot of samples prepare a 2 ml tube with 100 µl of $CaCl_2$ mix for each sample (wash buffer + $CaCl_2$ $C_F$ [2mM]) and resuspend the samples with it (before resuspension chill the tube in ice for 5')
- If you are using more than one PCR strip, add the $CaCl_2$ to each line with a 3' delay (needed for allowing the time to stop all the reactions later)

- If you are stopping the reactions using a multichannel pipette, you can leave 1/1:30’ in between the strips

6.3 Incubate in ice for 30’ sharp

6.4 Always in ice, place the PCR strip with the sample on the magnet rack, when the liquid is clear discard the supernatant.

Note: If the specific target has expected small fragments (e.g. 60 bp), don’t throw away the supernatant. Instead collect it, stop the reaction using EDTA and EGTA and merge it with the final elution later.

6.5 Resuspend each sample in 50 µl of 1X Urea STOP buffer

6.6 Incubate the sample 1h in the fridge 4°C (on a rotator)

6.7 Put the samples on the magnet rack and when the liquid is clear collect the supernanant and throw away the beads (collect the supernatant in new 200 µl PCR Tubes)

Note: the high concentration of the urea can, in time, denature the DNA, so after incubation proceed immediately to the beads clean up.

### 7 Bead clean up

For the clean-up use Mag-Bind®TotalPure NGS - M1378-01:

7.1 Warm up the beads to RT

7.2 In the meantime prepare a new aliquot (50 ml) of 80% EtOH

7.3 Prepare the Tris [10mM] pH8.2 (23°C)

7.4 Add 2X beads to each sample (100 µl beads for 50 µl sample)

7.5 Pipette up and down vigorously and multiple times to mix well (vortex can be used too); If doing this directly after elution of the fragments, you can start to prepare the tubes with the beads during the fragment release, in this way you can move the samples directly into the beads.

7.6 Incubate 15’ at room temperature

7.7 Put samples on the magnet rack and when the liquid is clear discard it

7.8 Without removing the tubes from the magnet rack or disturbing the beads wash the samples 2 times for 30" with 200 µl of 80% EtOH

Important! If you resuspend the beads, you will lose all the DNA because it is not bound to the beads

7.9 Remove the EtOH and dry (2-3’) the beads

7.10 Resuspend the beads in 25 µl of Tris [10 mM]

7.11 Incubate the samples 5’ at RT

7.12 Add 2x new beads and incubate 15’ RT (50 µl beads for 25 µl sample)

7.13 Without removing the tubes from the magnet rack or disturbing the beads wash the samples 2 times for 30" with 200 µl of 80% EtOH

7.14 Remove the EtOH and dry (2-3’) the beads

7.15 Resuspend the beads in 20 µl of Tris [10mM]

7.16 Incubate 5’ at RT

7.17 Put the sample on the magnet rack and when the liquid is clear move 20 µl of each sample in a new labeled tube

7.18 Proceed with library preparation

### 8 Library preparation:

To perform the library prep we are using the KAPA HyperPrep Kit (KR0961 - v.8.20).
Note! from this point best practice is to use only filter tips.

#### 8.1 End repair and A-Tailing:

- Perform the A-tailing following the instruction in the tables:

| Reagent | 1x |
| --- | --- |
| DNA<br>Buffer<br>Enzyme | 20µl<br>2.8µl<br>1.2µl |
| Total | 24µl |

| Thermocycler | Degree | Time |
| --- | --- | --- |
| A-Tailing | 12˚C<br>37˚C<br>58˚C | 15'<br>15'<br>45' |
| Hold | 4˚C | ∞ |

85˚C for the Lid

8.2 Adapter ligation:

8.2.1 Defrost the adaptors while centrifuging them at 4˚C 500 g 15' (If already in possession of the 1:10 dilution proceed from point 8.2.5)

8.2.2 Defrost the dilution buffer

8.2.3 In a 200 µl PCR tube take 2 µl of the right adaptor and 18 µl of dilution buffer (20 µl final volume)

8.2.4 Repeat the process using one tube for each adapter until you have a 1:10 stock of all the adapters needed

8.2.5 In a 200 µl PCR tube take 2 µl of the 1:10 dilution of the needed adapter and 18 µl of dilution buffer (obtaining a 1:100 dilution of the adapter)

8.2.6 Repeat the process using one tube for each adapter until you have a 1:100 stock of all the adapters needed (after using this dilution once trash the remaining)

8.2.7 Put the 1:10 dilution at -20˚C and use the dilution 1:100 (from next time start with this dilution and not from the original stock)

8.2.8 Prepare the reactions as follows:

| Reagent | 1x |
| --- | --- |
| Product | 24 µl |
| $H_2O$ | 2 µl |
| Buffer | 12 µl |
| Ligase | 4 µl |
| Adapters | 2 µl |
| Total | 44 µl |

| | | |
| --- | --- | --- |
| Ligation | 8˚C | Overnight |

8.2.9 Remember to write down sample and adaptors

8.3 Post ligation clean up (Mag-Bind®TotalPure NGS - M1378-01):

8.3.1 Warm up (RT) the beads for the clean up

8.3.2 In the meantime prepare a new aliquot (50 ml) of 80% EtOH from the 100% ultra pure bottle of EtOH

8.3.3 Prepare the Tris [10mM] pH8.2 (23°C)

8.3.4 In the same tubes used until now add 1.2X beads to each sample (50 µl beads for 44 µl sample)

8.3.5 Incubate 15’ at room temperature

8.3.6 Put on the magnet rack and when the liquid is clear discard it

8.3.7 Wash 2 times 30" with 200 µl of 80% EtOH without disturb the beads or removing the tubes from the magnet rack

8.3.8 Remove the EtOH and dry (not too much) the pellet

8.3.9 Resuspend the beads in 10 µl of Tris [10mM]

8.3.10 Put the sample on the magnet rack and when the liquid is clear move 10 µl of each sample in a new labeled PCR tube

8.4 Library amplification (13 cycles):

- Prepare the reaction as follow:

| Reagent | 1x |
|---|---|
| Library | 10 µl |
| Ready-mix | 12,5 µl |
| Primer-mix | 2.5 µl |
| Total | 25 µl |

| ID | 98°C | 45" |
|---|---|---|
| Denaturation | 98°C | 15" |
| Annealing | 60°C | 5" |
| Elongation | 60°C | 5" |
| Final elongation | 72°C | 1’ |
| Hold | 4°C | ∞ |

8.5 Sample clean up (Mag-Bind®TotalPure NGS - M1378-01):

8.5.1 Warm up (RT) the beads for the clean up

8.5.2 In the meantime prepare a new aliquot (50 ml) of 80% EtOH from the 100% ultra pure bottle of EtOH

8.5.3 Prepare the Tris [10mM] pH8.2 (23°C)

8.5.4 Without changing tube add 1.2X beads to each sample (30 µl beads for 25 µl sample)

8.5.5 Incubate 15’ at RT

8.5.6 Put on the magnet rack and when the liquid is clear discard it

8.5.7 Wash 2 time 30" with 200 µl of 80% EtOH without disturb the beads or removing the tubes from the magnet rack

8.5.8 Remove the EtOH and dry (not too much) the pellet

8.5.9 Resuspend the beads in 20 µl of Tris [10mM]

8.5.10 You can remove the beads right away or you can leave them there until the gel purification

8.6 Gel extraction (Using Thermo Fisher E-Gel):

After PCR amplification we load the product on Agarose 2% E-gels from Thermo Fisher. Each gel can have 10 sample + the ladder. We run the gel for 10’:

8.6.1 Take out the gel from the bag

8.6.2 Position the gel in the machine (electrode first) and press the gel until you hear a click

8.6.3 Only in case the clean up beads are still in the sample put the tubes on a magnet rack and wait until the liquid is clear, then proceed

8.6.4 Add to 20 µl of DNA sample to each well and 20 µl ladder in the first well (marked M)

8.6.5 Run 10’ with the right program taking picture every now and then

8.6.6 After the run, wait for the gel to cool down, open the gel chamber and cut out the DNA product in between the 150 bp and 500 bp (be sure to avoid the clear band at 120 bp, it is the adaptor dimers band)

8.6.7 Purify the DNA from the gel using a kit.

At this point you can proceed with sequencing
